# Residual cortical responses and network reorganization in inherited retinal degeneration: electrophysiological evidence

**DOI:** 10.64898/2026.08.18.745442

**Authors:** Ilaria Siviero, Alessia Verroca, Sonia Mele, Cora Miranda Lanza, Javier Sanchez Lopez, Chiara Quisisana, Valerio Marino, Silvia Francesca Storti, Leonardo Colombo, Daniele Dell’Orco, Paola Binda, Maria Concetta Morrone, Chiara Mazzi, Silvia Savazzi

## Abstract

Retinitis pigmentosa (RP) progressively deprives the retina of input, but whether the responsiveness of the visual cortex declines in parallel, remains preserved, or increases through compensatory gain remains unclear. Indeed, a weaker visually evoked response cannot, on its own, distinguish these possibilities, since it is equally compatible with a passively degraded input and with an actively recalibrated cortex. We combined spatially resolved steady-state visual evoked potentials (SSVEPs), which index stimulus-driven activity, with transcranial magnetic stimulation combined with electroencephalography (TMS-EEG), which probes cortical reactivity independently of vision, in patients with RP and in healthy controls.

Nine patients (PTs) with RP (five women, age range 28 to 69 years) and nineteen sex-, age-, and handedness-matched healthy controls (thirteen women, mean age 42.6 years) were tested. They underwent SSVEP recordings to stimuli presented at three eccentricities (central, intermediate, peripheral) and single-pulse TMS-EEG over the left and right occipital cortex and, as a non-visual control site, the dominant motor cortex. We quantified SSVEP amplitude and phase at 12 Hz, early TMS-evoked potentials, oscillatory power, inter-trial phase synchrony, and functional connectivity and graph-theoretical network measures derived from the weighted phase lag index.

SSVEP amplitude followed the expected central-to-peripheral gradient: PTs were comparable to healthy controls at the center, reduced but still above their own resting baseline at intermediate eccentricity, and no longer distinguishable from baseline in the periphery; phase differed from controls in a quarter of the central and half of the intermediate sectors. Occipital stimulation elicited a larger early negative deflection after left-hemisphere stimulation in PTs compared to controls, a stronger beta-band event-related spectral perturbation after stimulation of either hemisphere, and stronger, more efficiently distributed post-stimulus connectivity, despite comparable pre-stimulus connectivity, resting motor threshold, and most early evoked components. The pattern was site- and hemisphere-specific: left occipital stimulation produced widespread, mainly contralateral effects; right occipital stimulation a more circumscribed ipsilateral one, and motor cortex stimulation showed altered alpha-band activity without the bilateral occipital beta effect.

Together, these results show that progressive retinal deafferentation in RP does not produce a parallel decline in cortical responsiveness. Visually driven activity weakens with eccentricity, while direct cortical perturbation reveals preserved and, at selected sites, enhanced reactivity. This dissociation is consistent with a homeostatic increase in cortical gain rather than a uniform loss of cortical function, and indicates that the deafferented cortex retains, and in places strengthens, its capacity to respond as retinal input deteriorates.

## Introduction

Inherited retinal diseases (IRDs) comprise a genetically and phenotypically heterogeneous family of disorders in which retinal dysfunction or degeneration progressively compromises vision (Duncan et al., 2018). Retinitis pigmentosa (RP), the prototypical rod-cone dystrophy, is itself caused by mutations in many different genes but most often follows a recognizable functional sequence: rod dysfunction first impairs dark adaptation and peripheral vision, followed by progressive cone involvement and, at more advanced stages, loss of central vision (Verbakel et al., 2018). The resulting deprivation is therefore neither sudden nor spatially uniform. It advances across the retina over the years, leaving islands of residual sensitivity that vary markedly across patients and eccentricities. Although the primary pathology lies in the retina, its functional consequences extend to the entire visual pathway. This has become particularly important as restorative strategies move closer to clinical application: establishing whether a signal can still be generated at the retinal level is necessary, but it does not reveal how the visual cortex represents that signal, nor whether the cortex remains capable of responding to restored or newly introduced input.

Long-standing retinal deafferentation can alter both the structure and function of the visual brain. In RP, occipital gray matter reductions have been associated with the extent of visual field loss (Rita Machado et al., 2017), while resting-state studies have reported altered functional coupling within the visual system (Dan et al., 2019). Yet the nature of these changes remains unsettled. Responses within cortical territories deprived of their preferred retinal input have sometimes been interpreted as evidence of large-scale remapping. However, activity in these territories can depend strongly on the task performed by the observer, suggesting that feedback, attention, or the unmasking of pre-existing signals may contribute to the apparent recruitment (Masuda et al., 2010). Conversely, an eccentricity shift of central retinal representations toward more peripheral V1 locations has been described in patients with inherited peripheral retinal degeneration, in the absence of changes in V1 cortical thickness (Ferreira et al., 2017). Thus, the crucial question is not simply whether the deafferented visual cortex is active, but what drives that activity and what it reveals about the remaining network’s capacity for response.

Evidence from patients at different disease stages indicates that residual visual processing and cortical plasticity persist despite progressive retinal degeneration. Lunghi et al. (2019) showed that short-term monocular deprivation produced an ocular-dominance shift in RP comparable to the shift observed in healthy participants, and that the shift grew as retinal function deteriorated. In healthy adults, the same manipulation lowers resting GABA concentration in V1, and the size of that reduction predicts the perceptual shift (Lunghi et al., 2015), which places homeostatic regulation of the excitation and inhibition balance at the center of the mechanism. In animal models of RP, synaptic plasticity in the visual cortex is retained even when retinal degeneration is complete (Begenisic et al., 2020). In patients at a more advanced stage, Castaldi et al. (2019) found measurable sensitivity to low spatial and low temporal frequencies together with BOLD responses in striate and extrastriate cortex, even when standard electroretinographic and visual evoked responses were weak or absent and the flashes presented during scanning were not consciously perceived. Importantly, V1 responses tracked residual contrast sensitivity rather than cortical thickness. These findings argue against a visual cortex that becomes uniformly silent as retinal degeneration progresses, indicating instead that residual sensory processing and a capacity for homeostatic plasticity can persist in RP (Castaldi et al., 2020). They leave three mutually exclusive possibilities open: that cortical responsiveness declines in proportion to the loss of retinal drive, that it is preserved at a normal level, or that it increases as a stronger gain is applied to compensate for the incoming degraded visual signal. No measure of visually driven activity can separate them on its own, because a weaker visually driven response is compatible with all three once the input itself is degraded.

Electrophysiology is well-suited to this problem, although in RP it has been used mainly to characterize retinal output and conventional transient visual evoked potentials. Pattern-reversal visual evoked potentials (VEPs) remain recordable when pattern electroretinograms are already markedly abnormal (Janáky et al., 2008), which shows that cortical responses survive substantial retinal dysfunction. Steady-state visual evoked potentials (SSVEPs) offer a complementary measure with two separable components. Amplitude at the stimulation frequency indexes the strength of stimulus-driven activation, whereas phase indexes the temporal relationship between the stimulus and the entrained response (Norcia et al., 2015). The distinction matters here, because a reduction in the strength of the afferent signal and a delay in its arrival are dissociable: a phase shift without a loss of amplitude would point to altered timing of the afferent volley rather than to a smaller cortical response. Stäubli et al. (2026) recently showed that SSVEPs provide a reliable index of visual function requiring no behavioral response across the phenotypic spectrum of CRB1-related retinopathy, revealing both response attenuation and a shift of spatial-frequency tuning toward lower frequencies. Full-field or centrally weighted stimulation, however, cannot establish how cortical entrainment varies along the central-to-peripheral gradient that defines RP. The rationale for spatially selective stimulation is straightforward: only by driving discrete portions of the visual field separately can one test whether the amplitude and the timing of the cortical response follow the heterogeneous topography of residual vision.

SSVEPs, therefore, provide a spatially resolved measure of how residual visual input engages in the cortex. Complementing this measure of stimulus-driven activity, transcranial magnetic stimulation combined with electroencephalography (TMS-EEG) directly perturbs a cortical target and tracks both its local response and the subsequent spread of activity through connected regions with millisecond resolution (Ilmoniemi et al., 1997; Rogasch and Fitzgerald, 2013; Tremblay et al., 2019). Early TMS-evoked potentials provide an index of local cortical reactivity, whereas induced oscillatory activity, phase synchronization, and connectivity describe how the perturbation is expressed and propagated at the network level. Earlier TMS work in long-term pregeniculate blindness showed that increasing visual loss was associated with a lower probability and a more restricted spatial distribution of TMS-induced phosphenes, although phosphene thresholds remained within the normal range in patients who could still perceive them (Gothe et al., 2002). Building on this evidence, our TMS-EEG studies have shown that stimulation of the left and right occipital or parietal cortex engages distinct electrophysiological dynamics, with hemispheric asymmetries in local responses, signal propagation, functional connectivity, and directed effective connectivity (Siviero et al., 2023; Bonfanti et al., 2024; Paolini et al., 2024; Bonfanti et al., 2025). Any comparison of cortical reactivity between patients and controls must therefore be carried out separately for the two hemispheres, since collapsing across them would confound a group effect with the intrinsic asymmetry of the visual network.

In the present study, we used two complementary electrophysiological approaches in genetically characterized patients with RP and in sex-, age-, and handedness-matched healthy controls. Spatially selective SSVEPs were used to assess visually driven cortical responses, and TMS-EEG to probe cortical reactivity and signal propagation after stimulation of the left and the right occipital cortex, with stimulation of the dominant primary motor cortex as a non-visual reference. By combining a measure of stimulus-driven activity, resolved across the visual field in both amplitude and timing, with a direct perturbational measure of cortical responsiveness, we sought to determine whether the cortical consequences of retinal degeneration are best described by reduced responsiveness, by preserved responsiveness despite weakened afferent input, or by enhanced responsiveness consistent with homeostatic plasticity.

## Methods

### Section I - Population

Ten IRD patients (PTs) affected by RP and twenty sex-, age-, and handedness-matched healthy controls (HCs) participated in the study and underwent SSVEP and TMS-EEG sessions. However, one PTs and one HCs were excluded from the analyses due to low data quality. The analyses were conducted on nine PTs (5 females, mean age: 43.67, sd age: 13.79) and nineteen HCs (13 females, mean age: 42.63, sd age: 14.52). In Table 1, an overview of patients’ information is reported. In particular, one patient was left-handed, and so it was the corresponding age-matched HCs. The average symptom onset among patients was 20.44 years, while the symptom duration was 23.22 years. Data were collected in accordance with protocols approved by the local ethics committee (CARP - Comitato di Approvazione della Ricerca sulla Persona; 08.R1/2024). All subjects gave written informed consent in accordance with the Declaration of Helsinki.

**Table 1:** demographic and clinical information of IRD patients.

| ID | SEX | AGE | HANDEDNESS | GENE | SYMPTOMS ONSET (years) | SYMPTOMS DURATION (years) |
| --- | --- | --- | --- | --- | --- | --- |
| P01 | M | 32 | R | PRPH2 | 16 | 16 |
| P02 | M | 36 | R | RHO | 16 | 20 |
| P04 | M | 53 | R | RHO | 8 | 45 |
| P05 | F | 59 | R | RHO | 4 | 55 |
| P07 | F | 36 | R | RHO | 12 | 24 |
| P08 | F | 69 | R | RHO | 45 | 24 |
| P10 | F | 36 | L | PRPH2 | 18 | 18 |
| P11 | M | 44 | R | PRPH2 | 37 | 7 |
| P13 | F | 28 | R | RHO | 28 | 0 |

**Figure 1:**
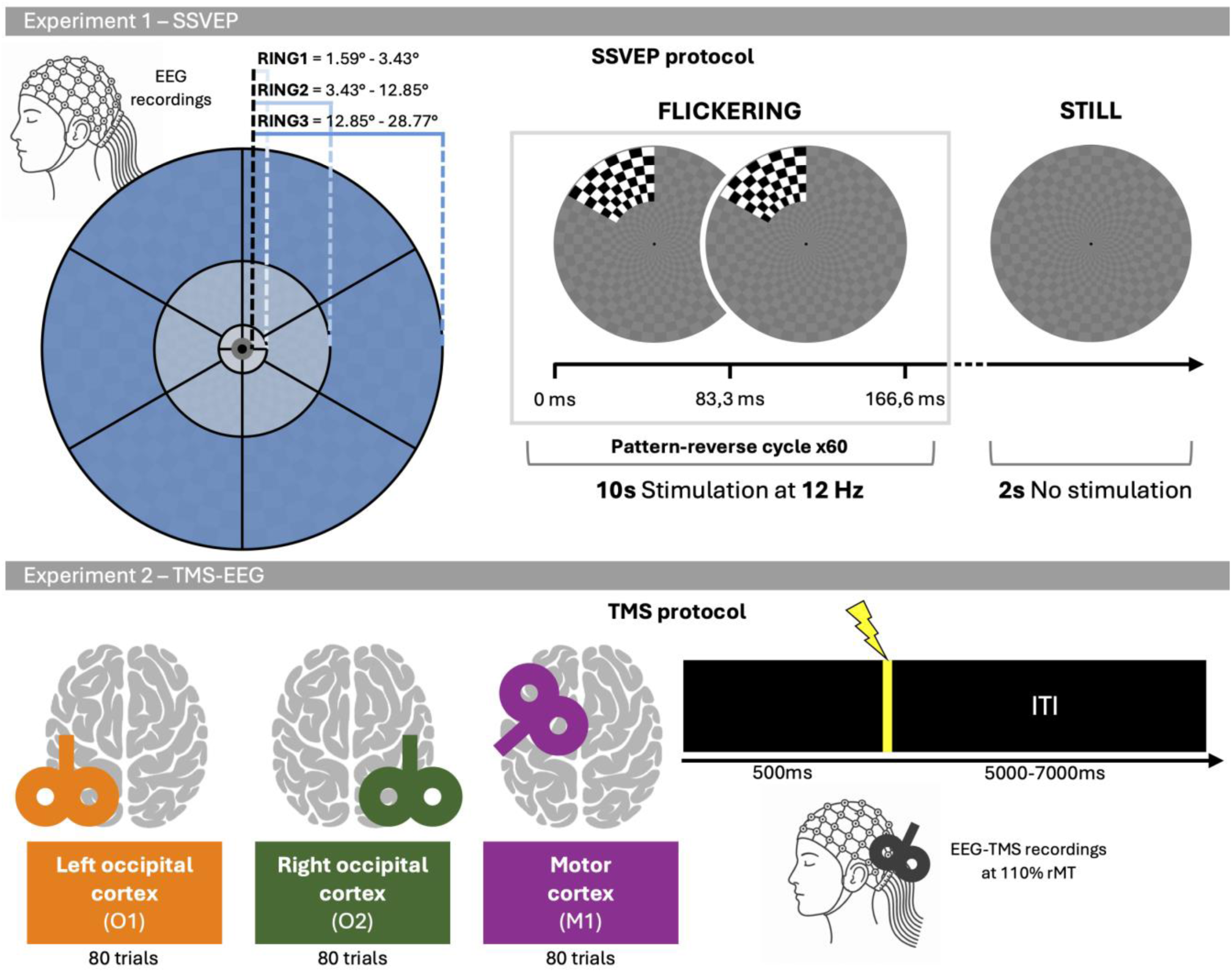
SSVEPs and TMS-EEG experimental settings and protocols.

### Section II – SSVEPs

#### SSVEP stimulation

Visual stimuli were generated with a custom-made Matlab script (R2024.b, Math-Works, Natick, MA, USA) developed for this study. The stimulus was a pattern-reversing checkerboard divided into 16 independently stimulable sectors, four of 90° in Ring 1 and six of 60° in Rings 2 and 3 (Capilla et al., 2016). Ring borders followed a geometric progression in shifted eccentricity coordinates, so that each ring activates approximately equal cortical territory according to the Rovamo–Virsu cortical magnification model (Rovamo & Virsu, 1979), which is more appropriate for scalp-level SSVEP than the purely striate model of Horton & Hoyt (1991). With E₂ = 10.22°, the ratio r = 1.690 gives borders at 3.43°, 12.85° and 28.77°, so that Ring 1 spans 1.59°–3.43°, Ring 2 spans 3.43°–12.85° and Ring 3 spans 12.85°–28.77°, for a stimulated field of 57.5°. The central 1.59° was never stimulated, to minimize contamination from fixation instability. Each sector contained 48 checks (8 angular by 6 radial), giving check sizes of 23.3, 77.6 and 161.3 arcmin at the mid-eccentricity of each ring (2.51°, 8.14° and 20.81°), corresponding to a spatial frequencies of 1.29, 0.39 and 0.19 cpd.

The screen background had a uniform luminance of 47.658 cd/m². The passive visual stimulation task consisted of two distinct, alternating conditions. In the STILL, or inter-stimulus, condition, a low-contrast checkerboard pattern was displayed, with checks alternating between 43.008 cd/m² and 51.145 cd/m²; in the FLICKERING condition, one pseudorandomly selected sector alternated between two complementary high-contrast black and white checkerboard frames (pattern and pattern reversal), with checks alternating between 0.233 cd/m² and 82.495 cd/m², while the remaining 15 sectors remained static in the low-contrast pattern. Each frame lasted 83.3 ms, corresponding to a frame rate of 12 Hz and an effective pattern-reversal frequency of 6 Hz: each trial comprised 10 s of FLICKERING followed by 2 s of STILL (12 s total), during which 120 frames (60 pattern-reversal cycles) were presented. A digital trigger, synchronized with the EEG acquisition, was sent at every frame onset (12 Hz). Stimuli were presented using E-Prime 3.0 (Psychology Software Tools, Pittsburgh, USA) on a 3840 x 2160 monitor at a viewing distance of 57 cm in a dark room.

Before the main experimental session, participants were shown a static black-and-white image of each of the 16 sectors and asked to report whether the sector appeared fully (1), partially (0,5), or not at all (0) visible. This preliminary assessment provided an in-session behavioral measure of binocular visibility of the stimulus as a function of eccentricity and polar angle, prior to the flickering stimulation blocks described below.

#### SSVEP procedure

Prior to the experimental blocks, two baseline conditions (open-eyes, OE; and closed-eyes, CE) were recorded, each lasting 2 minutes and including pseudo-triggers equivalent in number to the stimulation trials (1440 markers per condition). In these conditions, the screen continuously displayed a circular checkerboard with no flickering sectors, the same as the pattern used for the STILL condition. During the visual stimulation session, each of the 16 sectors was stimulated twice per block in pseudo-random order, with no consecutive repetitions of the same sector; each block comprised 32 FLICKERING trials (approximately 5.3 min), followed by a rest period, across a total of six blocks (192 trials; 12 repetitions per sector). Participants were instructed to fixate the central black dot throughout the session and to limit eye blinks and movements.

#### EEG Recordings

EEG data were recorded with a 59-electrode Ag/AgCl passive cap arranged according to the 10-10 International System (BrainAmp, Brain Products GmbH, Munich, Germany). The online reference was placed at the right mastoid (RM), while the ground electrode was at the AFz position. Horizontal and vertical eye movements were recorded using four electrodes placed at the left and right canthi, and at the upper and lower left positions of the eye. The impedance was kept below 5 kΩ throughout the session. For SSVEP stimulation, EEG data were recorded at a 1000 Hz sampling rate.

#### SSVEP data preprocessing and analysis

Raw EEG signals recorded during the SSVEP session were preprocessed with EEGLAB toolbox (version 2026.0; Delorme & Makeig, 2004). Firstly, a band-pass filter was applied between 0.01 and 49 Hz. The independent component analysis (ICA) was used to reduce artifacts related to eye blinks and movements. Their identification was supported by the ICLabel classifier (Pion-Tonachini et al., 2019). Then, continuous data were segmented into 2000 ms epochs time-locked to each pattern and pattern-reversal marker (i.e., every 83.3 ms, in line with the stimulation frame rate); consecutive epochs were therefore overlapped. Baseline correction was applied at the single-epoch level by subtracting its mean from each sample. Automated epoch rejection was based on multiple criteria: joint probability (epoch threshold: 5 sd; channel threshold: 3 sd), kurtosis (epoch: 5 sd; channel: 3 sd), amplitude threshold (±80 µV per channel), and linear trend detection (slope > 50 µV/s, R² > 0.3). Epochs with up to 6 problematic channels were corrected via spherical spline interpolation, whereas epochs exceeding this threshold were rejected; channels flagged in more than 30% of epochs were interpolated globally. These preprocessing steps were applied consistently across open-eyes baseline and stimulation conditions.

SSVEP data analyses addressed two complementary properties of the 12 Hz response: its amplitude and its phase. Amplitude quantifies the strength of the stimulus-driven activation, whereas phase indicates how this activation is time-locked to the stimulation, providing converging evidence that the response reflects genuine sensory entrainment rather than incidental activity at the same frequency.

Epochs were first averaged, and a Fast Fourier Transform (FFT) was then computed on the resulting waveform in order to assess both amplitude and phase of SSVEP data at 12 Hz. The FFT was applied to overlapping windows, each containing 2000 data points, with a frequency resolution of 0.5. Hz. This procedure isolates activity strictly phase-locked to the stimulation onset, since activity that is not time-locked to the marker is cancelled out by the averaging step (Vialatte et al., 2010; Norcia et al., 2015). Steady-state visual evoked responses were quantified as the SSVEP amplitude and phase at the stimulation frequency (12 Hz), averaged across an occipital region of interest (ROI: electrodes O1, O2, and Oz) where SSVEP responses were expected to be maximal (Norcia et al., 2015; Zhang et al., 2019). Circular average was used for phase data. For SSVEP amplitude, sector-level values were subsequently aggregated at the ring level (Ring 1: four sectors; Ring 2: six sectors; Ring 3: six sectors) by averaging 2000ms epochs. For SSVEP phase analysis, the sectors of the first two rings were assessed separately.

#### Statistical analyses of SSVEPs

For SSVEPs data, group differences in SSVEP amplitude at 12 Hz were assessed for each ring separately (R1, R2, R3) using two-tailed Wilcoxon rank-sum tests (HCs vs PTs, independent samples), with Bonferroni correction applied at ring-level comparisons. Effect sizes were expressed as the rank-biserial correlation (Kerby, 2014). Whole-ring amplitude within PTs and HCs was additionally compared against the baseline activity recorded during the open-eyes session. In this case, a one-tailed non-parametric test for paired samples was applied (Wilcoxon signed-rank test) and the Bonferroni correction was applied. This was applied to assess whether passive visual stimulation elicited neural activation that was both bigger than baseline (one-tailed hypothesis: SSVEP amplitudes at Ring 1, Ring 2, and Ring 3 exceeding the amplitude recorded during the open-eyes baseline condition) and specifically entrained with the stimulation frequency itself.

To assess group differences in phases extracted from the SSVEP waveform, Watson’s U² test was applied. This is a non-parametric test for unpaired circular data (Landler et al., 2025). Statistical significance was set at p < 0.05 throughout and the analyses were performed using Matlab (Version R2024b).

### Section III – TMS-EEG

#### TMS stimulation

The EEG signal was co-registered with the TMS and recorded at a sampling rate of 5000 Hz (see Section II for recording details). Coil positioning and targeting were based on an estimated individual MRI reconstructed from 3D head digitization obtained with Softaxic neuronavigation system (EMS, Bologna, Italy). During the stimulation, participants were placed in a dark room and fixated on a central gray cross displayed on a black background throughout the entire session; foam ear inserts were used to attenuate auditory-evoked responses to the TMS click sound.

Single-pulse TMS was delivered using a Magstim Rapid² stimulator (Magstim Company, Whitland, UK) equipped with a 70-mm figure-of-eight coil. TMS was delivered in a counterbalanced order over the left and right occipital visual cortex (localized using electrode positions O1 and O2, respectively) and over the primary motor cortex (identified by eliciting a twitch in the contralateral index finger) of the dominant hemisphere. One patient (P02) did not undergo TMS stimulation over the primary motor area. For occipital stimulation, the coil was placed tangentially to the scalp with the handle pointing upward and the coil plane-oriented perpendicular to the skull surface; for motor cortex stimulation, the coil handle was oriented downward at 45° from the midline. TMS stimuli were delivered at 110% of the resting motor threshold (rMT), which was determined in the dominant hand and defined as the minimum intensity required to elicit a motor evoked potential (MEP) ≥50 μV in at least 5 of 10 consecutive trials. For each site, the TMS pulses were divided into 4 blocks of 20 pulses each for a total of 80 trials.

#### EEG data preprocessing and analyses

EEG data co-registered during TMS were preprocessed using custom-made Matlab scripts (R2024.b, Math-Works, Natick, MA, USA) with EEGLAB toolbox (version 2026.0), and TMS–EEG signal analyzer (TESA) toolbox. The preprocessing steps followed the pipeline described in Rogasch et al. (Rogasch et al., 2017). The first step was to remove channels with excessive noise through visual inspection, followed by interpolation. Then, the continuous signals were segmented into 1000 ms epochs before and after the TMS pulse. The stimulation artifact was removed from −2 to 8 ms and replaced with the cubic interpolation method to avoid ringing artifacts. For cubic fitting, a 20ms time window before and after the removed data was used. Epoched data were down-sampled to 1000 Hz, and those contaminated by excessive artifacts were manually rejected. A first round of ICA was performed to remove TMS-related artifacts, and the signal from −2 to 8 ms was replaced with cubic interpolation. Data were filtered with a high-pass filter (>1Hz), followed by a low-pass filter (<70Hz) and with a band-stop filter (48–52 Hz). A second round of ICA was used to remove blinks, muscle activity, TMS-related decay artifact, and noisy channels. To improve component decomposition, the interpolated data from −2 to 8 ms after the TMS pulse were replaced with constant amplitude values before the ICA and interpolated again afterwards. The common average re-reference was applied, and data were baseline corrected considering the time window from -100 to -2 ms. Finally, data were down-sampled to 500 Hz. TMS-EEG data were analyzed through three steps: 1) early local component of evoked potentials, 2) time-frequency analysis of power and phase synchrony, and 3) phase-based brain connectivity and graphs.

##### 1) Early local components of evoked potentials

TMS-evoked potentials (TEPs) were calculated by averaging all trials for each participant in each stimulation condition. Then, the grand average was obtained by averaging TEPs across participants. For each condition, regions of interest (ROIs) were defined according to the stimulation site each comprising four electrodes surrounding the target area (left occipital: O1, Oz, PO3, PO7; right occipital: O2, Oz, PO4, PO8; motor cortex: C1, C3, CP1, CP3). For left-handed PTs and HCs, the EEG channels were flipped from right to left to align the two stimulation sites. Grand-average TEPs of the ROIs were visually inspected separately for HCs and PTs to identify the first positive and first negative components. A common time window of interest was selected based on HCs and subsequently applied to both groups. Of note, in most cases, peak latency was the same in both groups and differed by no more than 2 ms. Specifically, a time window of 12 ms (±6 ms around the peak) was centered respectively on the first positive and the first negative peak identified in each ROI for both groups. The selected time windows were as follows: left occipital ROI, P20 (14–26 ms) and N35 (28–40 ms), right occipital ROI, P20 (16–28 ms) and N35 (30–42 ms), and motor cortex ROI, P30 (24–36 ms) and N45 (36–48 ms). Therefore, for each participant, TEP amplitude was quantified as the mean amplitude over the time window of 12 ms within the corresponding ROI. This analysis allowed us to evaluate group differences in the earliest local TEP (within 50 ms after TMS) around the target area, which are indices of cortical excitability.

##### 2) Time-frequency power and phase synchrony

In this study, a complex Morlet wavelet ranging between 8 and 45 Hz (frequency resolution = 0.5 Hz, number of cycles = 3.5) (Dworkin et al., 2025) was applied using the open toolbox presented by Morales et al. (Morales et al., 2022). TMS-induced oscillatory power was then quantified through the event-related spectral perturbation (ERSP), which reflects changes in total power relative to the pre-stimulus baseline, including both phase-locked and non-phase-locked activity. Therefore, the time-frequency spectral estimates were normalized with the baseline interval using the divisive method (Morales et al., 2022). The time window from −500 to −100 ms was selected as baseline, and subsequently ERSP values were expressed in decibel (log-transformation). Finally, the global ERSP was obtained by averaging the ERSP over all channels.

Phase consistency across trials was quantified by considering the phase angle at each time and frequency point. This measure is commonly referred to as inter-trial phase synchrony (ITPS; but also, as inter-trial phase coherence [ITPC]). It ranges between 0 (no phase consistency) and 1 (perfect phase consistency).

Both ERSP and ITPS were separated into three frequency bands (alpha [8–13 Hz], beta [13–30 Hz], and gamma [31–45 Hz]) and averaged within 20-400 ms (Vallesi et al., 2021).

##### 3) Brain functional connectivity and graph network analysis

Functional connectivity was calculated using the weighted phase lag index (wPLI). It is a well-known measure in the field of TMS-EEG that allows assessment of the non-zero lag-phase difference between pairs of electrodes (De Martino et al., 2024) wPLI is an extension of the phase lag index (PLI), with the advantage of accounting for volume conduction and reducing the risk of missing true instantaneous connections. In this study, the wPLI was computed across trials to track brain connectivity consistency across TMS pulses. A seed-based analysis was performed to evaluate how stimulation caused a phase shift in the brain network. For each stimulation site, O1, O2, and C3 electrodes were selected as seed channels, respectively, and their connectivity with all other electrodes was evaluated. The C4 electrode was used for the patient and healthy control stimulated over the right motor cortex. wPLI values were then averaged in the time window between 20 and 200 ms in order to investigate the effect of TMS pulses within an early time window after stimuli. Also, the pre-stimulus time window (−500 to −100 ms) was considered to investigate the differences between the two groups.

To investigate brain re-organization and/or alteration in IRD patients with respect to HCs, graph analysis was employed to evaluate both segregation and integration properties of the brain networks (De Vico Fallani et al., 2014). In particular, segregation reflects the brain’s ability to process local information within a brain region or its interconnected groups. On the other hand, the property of integration concerns the ability to combine information from different brain regions, e.g., distant ones. In our context, these could allow assessing network-level plasticity triggered by progressive sensory deprivation.

For each participant, strength, global and local efficiency, clustering, characteristic path length and small-world propensity (SWP) were calculated for their corresponding wPLI networks. In a brain network, each electrode is represented as a node, while the relationships between two electrodes correspond to links. Node strength is defined as the sum of the weights of all links connected to a given node. Global efficiency measures the average inverse shortest-path length across the network and is inversely related to the characteristic path length. Local efficiency is calculated as the global efficiency of a node’s neighborhood and is closely related to the clustering coefficient. Finally, SWP quantifies the extent to which a network displays small-world structure, and it is measured as the deviation of a network clustering coefficient and characteristic path length from both regular and random graphs. These measures are typically used to characterize the topological organization of brain networks and to identify alterations relative to healthy controls or within the same population across different conditions/time points (Zhang et al., 2025).

As previously described, the graph metrics were calculated over 20-200 ms for each frequency band. Due to the variability of the brain connectivity within and between subjects, multiple thresholds were applied to the networks to compare the two populations and different TMS-conditions. Sparsity thresholds ranged from 0.3 to 0.5 with a step of 0.05 (i.e., 0.3 refers to 30 percent of the strongest connections are maintained). Graph measures were calculated for the above threshold range using the brain connectivity toolbox (BCT). Finally, to assess differences between PTs and HCs, the area under the curve (AUC) was estimated for each metric (De Vico Fallani et al., 2014).

### Statistical analyses of TMS-EEG data

The mean TEP amplitudes within each time window of interest, as well as ERSPs and ITPS within each frequency band, were compared between groups using the Wilcoxon rank-sum test. For each stimulation site, a Bonferroni correction was applied to account for multiple comparisons (*p<*0.05). Differences in wPLI were assessed through the cluster-based permutation test, as implemented in FieldTrip toolbox (*p<*0.05). Finally, group differences in the AUC extracted from each graph measure were assessed using the Wilcoxon rank-sum test (p < 0.05).

## Results

### Section I – Clinical and demographic data analysis

No statistically significant differences in age between the two groups were observed (U = 90.50, z = 0.224, p = 0.8227, rbc = 0.058). Figure 2 shows Humphrey Visual Field (HVF) test results for the left (LE) and right (RE) eye of each patient. HVF assessed with Humphrey Visual Field Analyzer 30-2 SITA-Fast perimetry protocol (Bengtsson & Heijl, 1998). Overall, the visual field pattern was heterogeneous across patients. In P04, P05 (RE only), P07, P08, and P10, visual loss was extensive, with only a small residual island of preserved sensitivity in the center surrounded by an otherwise non-accessible periphery, consistent with the peripheral constriction and central island preservation typically described in IRD (Hamel, 2006). In P01, P02, P11, and P13, larger portions of the visual field retained residual, though patchy and non-uniform, visual field access.

For the TMS session, the mean rMT of HCs was 71.42, and the sd was 11.82, while the mean for PTs was 69.78, and the sd was 10.92. The rMTs did not differ statistically between PTs and HCs (U = 83.50, z = -0.074, p = 0.9411, rbc = -0.023).

**Figure 2:**
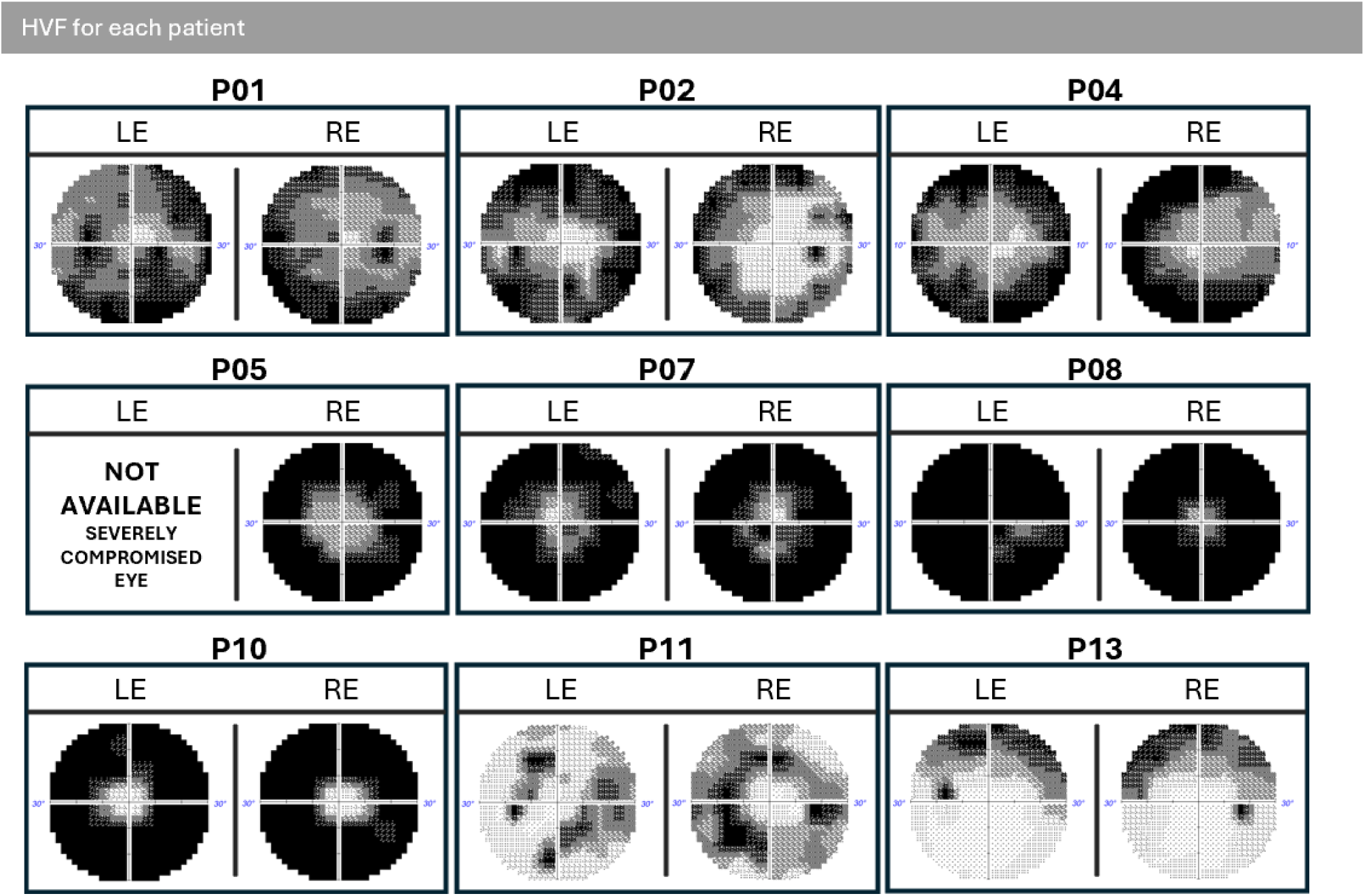
In the grayscale plots, darker regions correspond to areas of reduced or absent light sensitivity, while lighter regions correspond to areas of preserved sensitivity; the four quadrants are centered on fixation, with eccentricity extending to 30°. To note, due to severely compromise vision, perimetry could not be performed on the left eye of P05, and only 10° of the visual field for each eye was assessed for P04.

### Section II – SSVEP results

**Figure 3:**
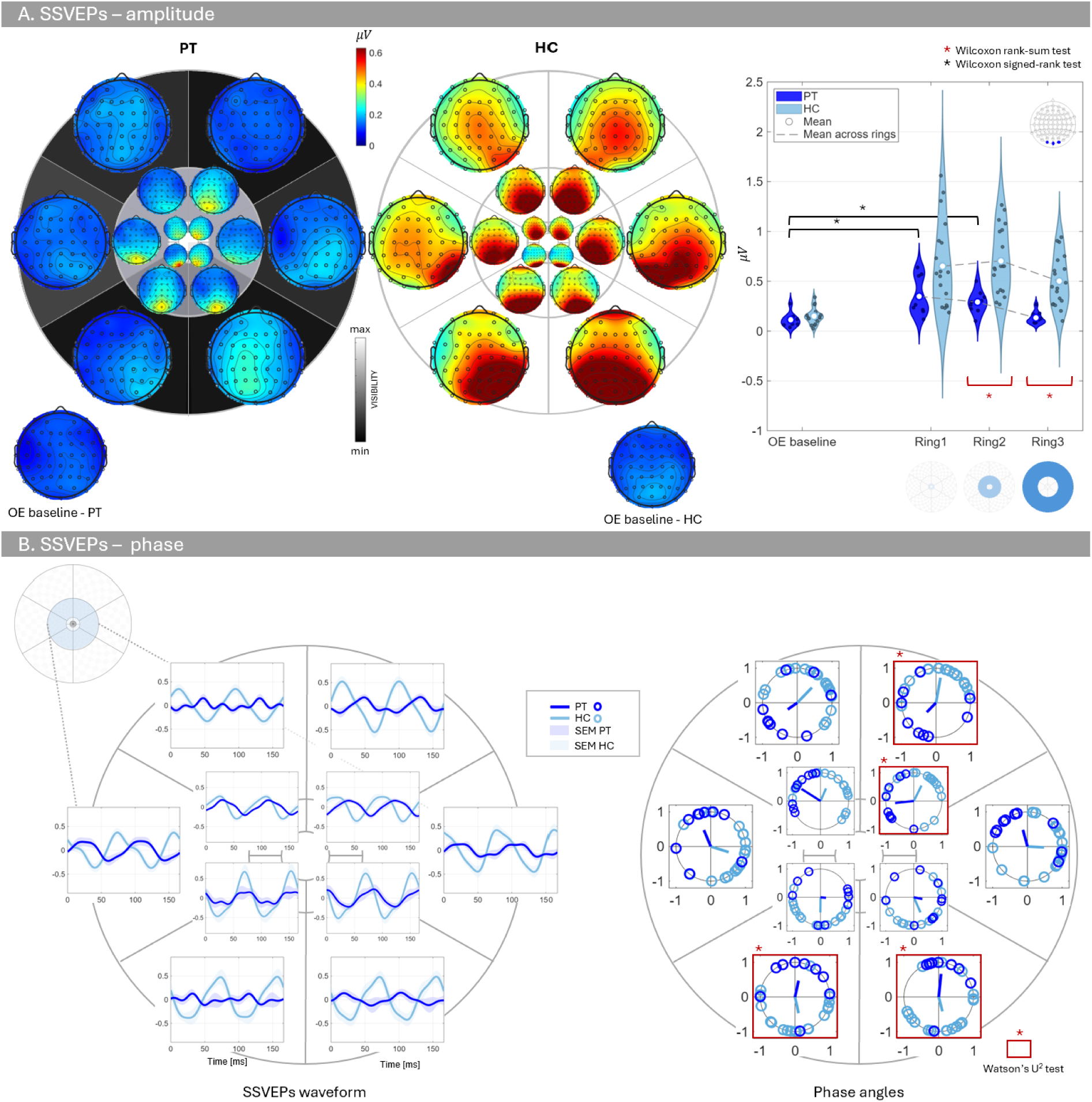
SSVEP amplitude and phase results. **A)** Left: Grand-average scalp topographies of the mean SSVEP amplitude at 12 Hz for the EO and flickering conditions, shown separately for PTs (left) and HCs (middle). Sector-level visibility was represented on a grayscale ranging from black (lowest mean visibility across participants) to white (highest mean visibility, corresponding to all participants reporting full visibility of that sector). Accordingly, the background shading of each sector indicates its mean subjective visibility (fully, partially, or not visible) across the two groups of participants, as assessed prior to the experimental session (see Methods). Right: statistical analysis of SSVEP amplitude. Mean SSVEP amplitude (ROI: O1, O2, and Oz) for each of the 3 stimulated rings for PTs and HCs in baseline open-eyes and visual stimulation conditions. The amplitude in each ring was compared using non-parametric tests against the open-eyes baseline (PTs vs OE baseline) and flickering condition (PTs vs HCs) with Bonferroni correction (*p<*0.05). **B)** Left: mean waveform of SSVEP with standard error of mean (SEM) over O1, O2 and Oz for the first and the second ring. Right: phase representation and statistical analysis.

To assess whether ring-level SSVEP amplitude reflected genuine stimulus-driven entrainment rather than a difference already present at rest, amplitude within PTs was compared against a within-subject baseline open-eyes condition using one-tailed Wilcoxon signed-rank tests, with Bonferroni correction. SSVEP amplitude in PTs was significantly higher than open-eyes baseline in Ring 1 (W = 42, p = 0.030, rbc = 0.867, Bonferroni corrected) and Ring 2 (W = 43, p = 0.018, rbc = 0.911, Bonferroni corrected) but did not differ from baseline in Ring 3 (W = 29, p = 0.744, rbc = 0.289, Bonferroni corrected). This result indicates that, only in the most peripheral ring, PTs failed to show activation above its own resting level.

The same comparison was carried out in HCs. Amplitude in HCs was significantly higher than open-eyes baseline in Ring 1, Ring 2, and Ring 3 (all: W = 190, z = 3.803, p < 0.001, rbc = 1.000, Bonferroni corrected), indicating that stimulus-driven activation exceeded resting baseline at every eccentricity. This contrasts with PTs, in whom activation above baseline was confined to Ring 1 and Ring 2, and was absent in the most peripheral ring. Group differences in SSVEP amplitude at 12 Hz were assessed for each ring separately using two-tailed Wilcoxon rank-sum tests (Figure 3A). PTs showed lower amplitude than HCs in Ring 2 (U = 16, z = - 3.39, p = 0.003, rbc = - 0.813, Bonferroni corrected) and Ring 3 (U = 7, z = -3.84, p < 0.001, rbc = -0.918, Bonferroni corrected). In Ring 1, PTs and HCs amplitude differences were not statistically significant (U = 49, z = -1.77, p = 0.306, rbc = -0.427, Bonferroni corrected). In addition, HCs and PTs were compared on the baseline open-eyes condition to verify that the group difference did not already exist independently of the visual stimulation; this comparison confirmed quantitatively what was indicated qualitatively above, with no significant difference between groups (U = 56, z = -1.43, p =0.615, rbc = -0.345, Bonferroni corrected). Finally, SSVEP results revealed a reliable stimulus-driven cortical response at 12 Hz in both groups, with an amplitude gradient modulated by eccentricity that was reduced in patients relative to healthy controls, particularly over the more peripheral portions of the stimulated visual field.

Phase differences were assessed only for Ring 1 and Ring 2, where PTs elicited a neural response, as confirmed by SSVEP amplitude analysis. For this reason, in the left part of Figure 3B, the phase angles are reported for the first two rings. Each plot shows the angle for each participant (PTs and HCs), separated for each sector. The direction of the colored arrow represents the circular mean phase of each group, while its length is proportional to the mean resultant vector length (R), providing a measure of phase concentration. Phase angles were statistically different between PTs and HCs in 25% of the sectors in Ring 1 (p = 0.006, Bonferroni corrected) and 50% of the sectors in Ring 2 (p=0.037, p = 0.032, and *p<<*0.05, Bonferroni corrected). Although SSVEP amplitude did not differ significantly in Ring 1, the phase presented a shift of 122.69 degrees (i.e., delta_phases = PTs-HCs). In Ring 2, significant phase shifts of 155.49, 159.55, and 145.65 degrees were observed in the lower sectors (left and right) and the upper-right sector, respectively. Compared with Ring 1, Ring 2 exhibited larger differences with the circular mean phase of PTs being consistently shifted relative to HCs across the significant sectors.

### Section III – TMS-EEG results

**Figure 4:**
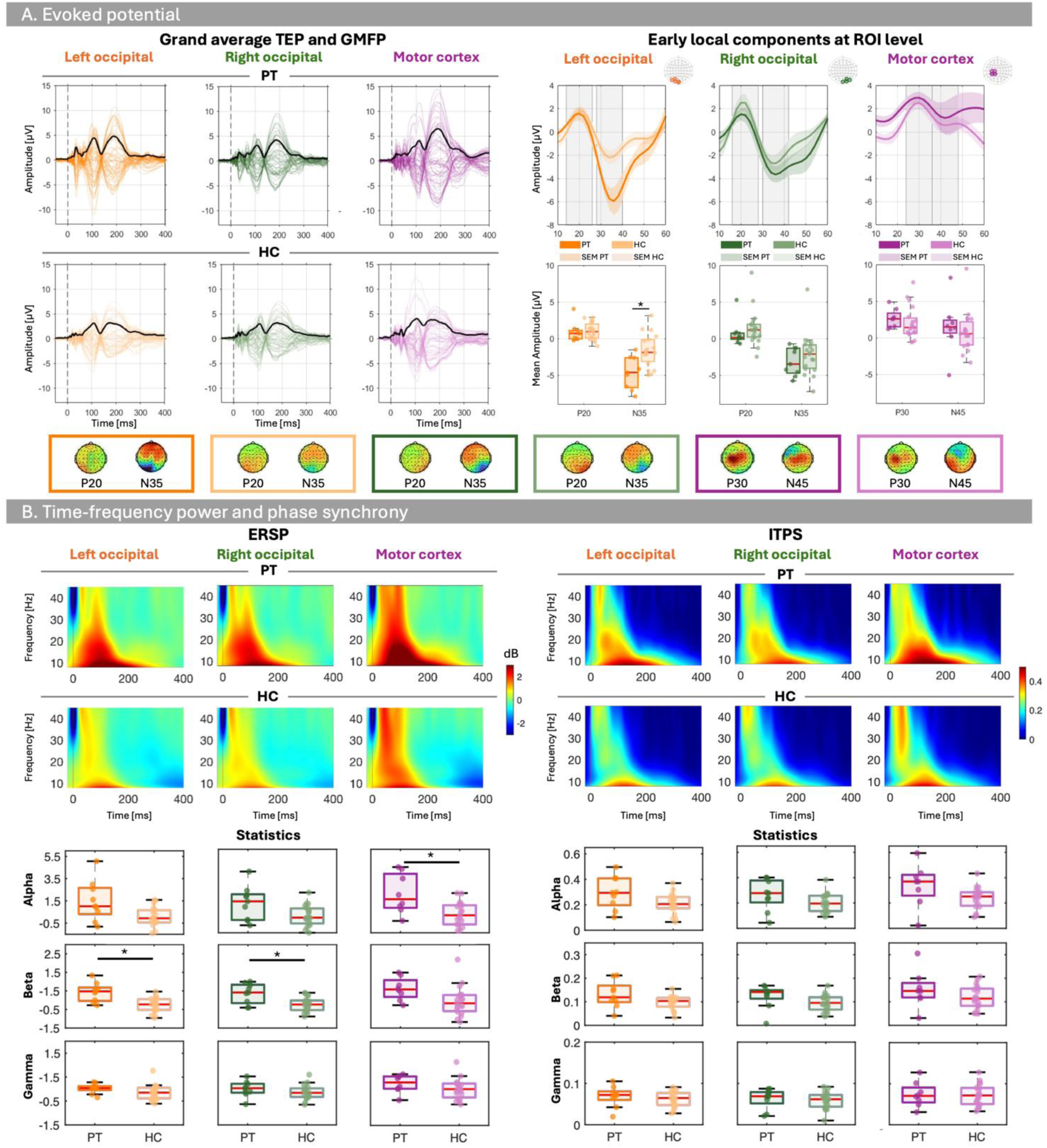
Evoked potential and time-frequency results. **A)** Comparison of TEPs between PTs and HCs. The left panels show the grand-average TEP waveforms elicited by stimulation of the left and right occipital cortex, and motor cortex (from left to right) in PTs (upper row) and HCs (lower row), with the black line representing the global mean field power (GMFP). Scalp topographies illustrate the analyzed TEP components for each positive and negative peak within the corresponding region of interest (ROI). In the upper row of the right panels are represented the grand-average TEP waveforms extracted from the ROI defined for each stimulation site. The gray shaded area indicates the time window surrounding the peak of interest. Under them, boxplots compare the mean peak amplitude between PTs and HCs. **B)** Time–frequency analyses of power and phase. The left panel shows the ERSP results for each stimulation site: left occipital (O1, orange), right occipital (O2, green), and motor cortex (M1, purple). Time–frequency representations are displayed for the PTs (upper row) and HCs (lower row). The lower panel presents boxplots of ERSP values for each frequency band and stimulation site in PTs and HCs. The right panel shows the corresponding ITPS results. Statistically significant group differences are indicated by asterisks (Bonferroni-corrected, *p<*0.05).

TMS-EEG results revealed altered evoked activity in IRD patients following visual cortex stimulation, pointing to reorganization within the affected cortical circuits. TEP components showed similar timing and spatial distribution between IRD patients and HCs. However, the left occipital N35 negativity was visibly enhanced in IRD patients (mean –4.645; sd 2.191) compared to HCs (mean –1.635; sd 2.211). The analysis highlighted a significant difference in the mean amplitude of the first negative component (N35) following left occipital stimulation between the PTs and HCs groups (U = 28, z = -2.804, p = 0.005, rbc = -0.672). No significant group differences were observed for the mean amplitude of any other early component.

As shown in Figure 4B, significant differences in ERSPs between PTs and HCs were observed across the full 20-400 ms post-stimulation window. In particular, in the case of left and right occipital stimulation (i.e. O1 and O2), group differences were found in the beta band (U = 148, z = 3.001, p = 0.003, rbc = 0.719; U = 138, z = 2.558, p = 0.010, rbc = 0.614, Bonferroni corrected). Conversely, differences in the alpha band were found following the motor cortex stimulation (U = 122, z = 2.416, p = 0.016, rbc = 0.605, Bonferroni corrected). Particularly, PTs showed stronger ERSPs than HCs, highlighting how stimulation causes global activation throughout the analyzed period in patients. On the other hand, the analysis conducted on ITPS did not reveal any statistically significant differences. This suggests that PTs may exhibit unchanged brain activity in phase synchrony across the entire time interval compared with HCs. Moreover, it is interesting to note that there is no difference in phase synchronization latency between the two groups.

**Figure 5:**
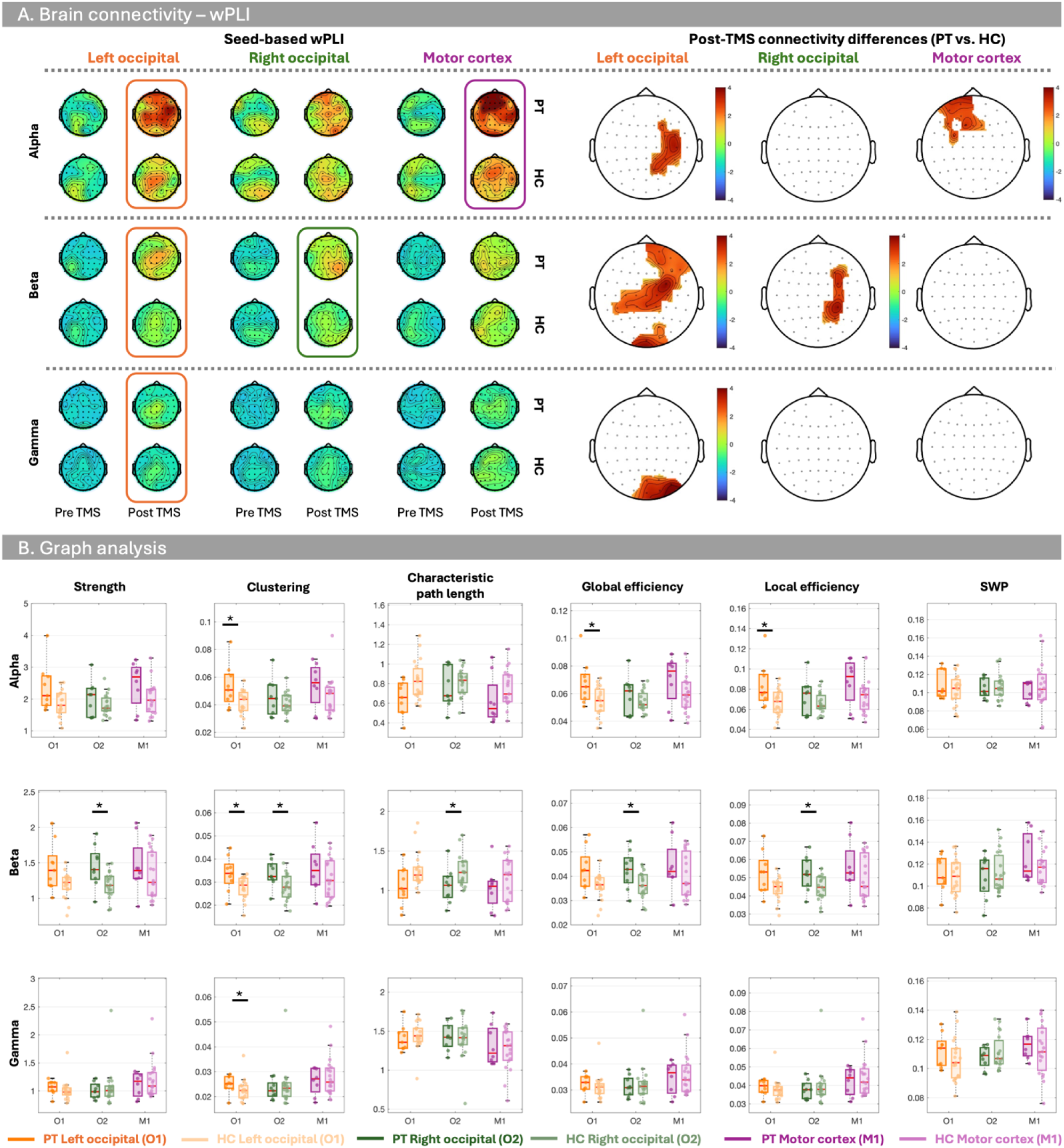
Brain connectivity and graph analysis. **A)** Functional connectivity estimated using the wPLI in the alpha, beta, and gamma frequency bands following stimulation of the left occipital cortex (O1), right occipital cortex (O2), and motor cortex (M1). Topographic maps represent the seed-based wpli values for PTs and HCs. Cluster-based permutation test highlights statistically significant differences between groups, with positive (red) values indicating higher connectivity in PTs than HCs. **B)** Group comparison of graph metrics derived from the weighted connectivity matrices, including network strength, clustering coefficient, characteristic path length, global efficiency, local efficiency, and SWP, for each stimulation site and frequency band. Each point is the AUC under the selected sparsity thresholds. Asterisks refers to statistically significant differences between groups (*p* < 0.05).

In Figure 5A, the brain connectivity results are shown. On the left part, for each frequency band, topographies of the seed-based wPLI values are shown both in the pre-TMS period and within 20-200 ms. Of particular interest, comparable values across groups within each frequency band were observed during the pre-TMS period, suggesting that these connections may remain unchanged relative to HCs. On the other hand, we observed a stronger and broader post-TMS hyper-synchronization in PTs, especially in the alpha band.

These suggest that a compensatory hyper-responsiveness could appear in IRDs patients, at the same time with a systemic altered expression and propagation of the TMS-evoked response involving both visual and non-visual areas. As to the cluster-based permutation tests showed in the right part of Figure 5A, statistically significant differences were found in: 1) alpha band for the left occipital stimulation and the motor cortex stimulation; 2) beta band following the left and the right occipital stimulation and 3) gamma band following the left occipital stimulation. In particular, targeting the left occipital brain areas revealed contralateral, widespread differences between PTs and HCs across the brain relative to the site of stimulation. On the other hand, by targeting the right occipital site, the differences were circumscribed to ipsilateral channels, especially parietal and central channels.

Figure 5B shows the results of graph-based analysis for each stimulation site and frequency band. Changes in topological networks between PTs and HCs were observed following left occipital stimulation for clustering coefficient across all frequency bands, and for global and local efficiency in the alpha band. Following right occipital stimulation, significant group differences were found in network strength, clustering coefficient, characteristic path length, and global and local efficiency in the beta band. No significant differences were observed in SWP for any stimulation site and frequency band. Overall, increases in strength, clustering coefficient, and global and local efficiency were observed for PTs relative to HCs, while a reduction in characteristic path length was observed for PTs. Table 2 presents complete statistical results for the graph measures.

**Table 2:**
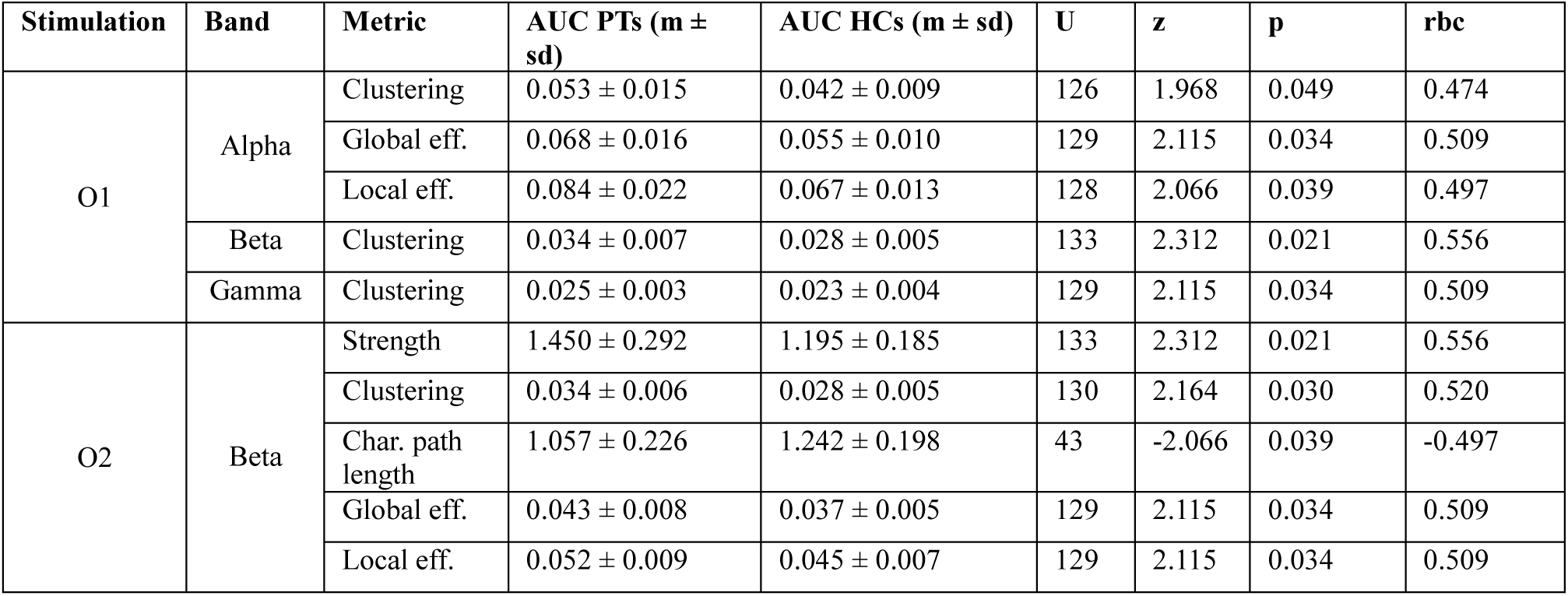
statistically significant results (*p<*0.05) of graph measures.

| Stimulation | Band | Metric | AUC PTs (m ± sd) | AUC HCs (m ± sd) | U | z | p | rbc |
| --- | --- | --- | --- | --- | --- | --- | --- | --- |
| O1 | Alpha | Clustering | 0.053 ± 0.015 | 0.042 ± 0.009 | 126 | 1.968 | 0.049 | 0.474 |
| | | Global eff. | 0.068 ± 0.016 | 0.055 ± 0.010 | 129 | 2.115 | 0.034 | 0.509 |
| | | Local eff. | 0.084 ± 0.022 | 0.067 ± 0.013 | 128 | 2.066 | 0.039 | 0.497 |
| | Beta | Clustering | 0.034 ± 0.007 | 0.028 ± 0.005 | 133 | 2.312 | 0.021 | 0.556 |
| | Gamma | Clustering | 0.025 ± 0.003 | 0.023 ± 0.004 | 129 | 2.115 | 0.034 | 0.509 |
| O2 | Beta | Strength | 1.450 ± 0.292 | 1.195 ± 0.185 | 133 | 2.312 | 0.021 | 0.556 |
| | | Clustering | 0.034 ± 0.006 | 0.028 ± 0.005 | 130 | 2.164 | 0.030 | 0.520 |
| | | Char. path length | 1.057 ± 0.226 | 1.242 ± 0.198 | 43 | -2.066 | 0.039 | -0.497 |
| | | Global eff. | 0.043 ± 0.008 | 0.037 ± 0.005 | 129 | 2.115 | 0.034 | 0.509 |
| | | Local eff. | 0.052 ± 0.009 | 0.045 ± 0.007 | 129 | 2.115 | 0.034 | 0.509 |

## Discussion

In the present study, we tested nine patients with RP and nineteen matched healthy controls with two complementary techniques. We used spatially resolved SSVEPs to test whether the strength and timing of stimulus-driven cortical responses follow the central-to-peripheral pattern of visual loss that characterizes RP. Moreover, we used TMS-EEG to test, independently of any visual stimulus, how the same patients’ cortex responds when directly perturbed. With SSVEPs, we found that response amplitude was comparable to controls in the central ring, reduced but still above the patients’ own resting baseline in the intermediate ring, and no longer distinguishable from baseline in the periphery; phase was also altered in several sectors of the two rings that retained a measurable response. With TMS-EEG, we found the opposite pattern: occipital stimulation elicited an enhanced early N35 after left-hemisphere stimulation, a stronger beta-band ERSP after stimulation of either hemisphere, and a stronger and more widely distributed post-TMS connectivity, despite comparable resting motor thresholds and comparable pre-TMS connectivity. These results indicate that retinal deafferentation in RP does not produce a parallel weakening of cortical responsiveness: visually driven activity becomes progressively weaker with eccentricity, while the same cortex, when perturbed directly, responds as strongly as, or more strongly than, that of healthy controls.

The SSVEP results extend previous reports of attenuated steady-state responses in inherited retinal disease (Stäubli et al., 2026) by showing that attenuation is not uniform across the visual field but follows the topography of residual vision shown by the RP patients. Open-eyes baseline amplitude did not differ between groups, and healthy controls showed activation above baseline in all three rings, so the absent response in the patients’ periphery reflects a genuine loss of stimulus-driven entrainment, not a group difference in ongoing activity at the stimulation frequency. Conversely, the response above baseline that we still recorded in Ring 2 shows that reduced amplitude relative to controls reflects diminished, not abolished, cortical entrainment. Thus, ring-level SSVEP amplitude captured a graded loss of stimulus-brain coupling that mirrors, at the cortical level, the central island of preserved sensitivity typically found on perimetry in RP. Moreover, phase differed between groups in a quarter of the sectors in Ring 1 and in half of the sectors in Ring 2, with shifts of the circular mean phase of about 123 degrees in Ring 1 and between 146 and 160 degrees in the affected Ring 2 sectors; phase could not be assessed in Ring 3, where no stimulus-driven response survived above baseline. A shift of this size in Ring 1, where amplitude itself did not differ from controls, shows that the temporal relationship between stimulus and cortical response can be altered in RP before amplitude itself is affected.

As for TMS-EEG, the results argue against a simple, generalized increase in cortical excitability. Earlier work in long-standing pregeniculate blindness (Gothe et al., 2002) had already suggested that the relationship between visual loss and cortical excitability is not linear: the probability and spread of TMS-induced phosphenes fell as blindness became more severe, yet phosphene thresholds stayed within the normal range as long as some vision remained. If retinal deafferentation in RP produced an undifferentiated increase in cortical excitability, we would expect it to appear regardless of hemisphere, frequency band, or component. Instead, the N35 enhancement was confined to left occipital stimulation; beta-band ERSP was increased after stimulation of either occipital site, but ERSP increased without a corresponding increase in inter-trial phase synchrony in the whole-time window (20-400ms), so the perturbation generated more oscillatory power without a stronger or more consistent phase reset across trials; resting motor threshold and most early TEP components did not differ between groups. Importantly, this selective, site- and measure-dependent pattern is what a system-level recalibration of gain, rather than a global loosening of inhibition, would predict.

Beyond this local response, the TMS-EEG connectivity results locate this enhanced response within the network rather than at a single site. Before the TMS pulse, wPLI values were comparable between patients and controls; the difference emerged only after the pulse, together with increased clustering and local efficiency, in keeping with stronger coupling among neighboring nodes, and increased global efficiency with a shorter characteristic path length, in keeping with perturbation-related activity reaching more distant nodes through shorter functional paths. Small-world propensity, in contrast, did not differ, so the overall balance between segregation and integration was preserved even as its underlying strength and efficiency changed. Because graph measures are mathematically interdependent, we do not treat each metric as an independent line of evidence. Together, however, they point in the same direction: after TMS, patients’ networks couple more strongly among neighboring nodes and distribute the perturbation more widely than controls’ networks do. Thus, a single pulse delivered at one node propagates across a larger portion of the network, and does so more efficiently, in patients than in controls.

A homeostatic increase in cortical gain (Castaldi et al., 2020) provides a single, parsimonious account of this dissociation, preferable to positing two independent processes, one weakening visually driven entrainment and another, unrelated, enhancing perturbational responsiveness. Indeed, the same retinal degeneration that impoverishes visual drive could also be the trigger that raises cortical gain, so that a weaker entrained response and an enhanced perturbational response become two expressions of one underlying change rather than two separate phenomena requiring two separate explanations. The same logic extends to the SSVEP gradient: if gain acts on whatever attenuated afferent signal still reaches the cortex, amplitude in Ring 1 and Ring 2 need not fall as far below control values as the underlying retinal loss alone would predict, which is consistent with amplitude in Ring 1 not differing from controls and remaining above the patients’ own baseline in Ring 2. No degree of cortical amplification, however, can generate an entrained response from an input that no longer arrives, which is what we take in the absence of any response above baseline in Ring 3 to reflect. Weaker SSVEPs and enhanced TMS-evoked responses are, on this view, two windows onto the same compensatory process; one still bounded by how much retinal signal survives, the other largely independent of it. This interpretation sits within a body of evidence, largely from Morrone and colleagues, that the adult visual cortex retains, and in RP may even upregulate its capacity for homeostatic plasticity. For instance, ocular-dominance plasticity elicited by short-term monocular deprivation is preserved in RP and grows stronger, not weaker, as retinal function deteriorates (Lunghi et al., 2019). In healthy observers, the same manipulation lowers V1 GABA, and the size of that reduction predicts the size of the perceptual shift (Lunghi et al., 2015), which ties ocular-dominance plasticity directly to a reduction in cortical inhibition. Moreover, residual visual signals can still drive V1 and extrastriate BOLD responses at advanced stages of retinal disease, and V1 activity tracks residual contrast sensitivity rather than cortical thickness (Castaldi et al., 2019), which shows that the deafferented cortex remains functionally coupled to whatever input still reaches it. Finally, preserved visual-cortical plasticity in animal models of retinal degeneration provides converging support from different species and a different method (Begenisic et al., 2020). Together, these studies converge on the claim our results extend: the visual cortex retains, and in RP may redeploy, an active capacity to recalibrate its own responsiveness as retinal input degrades.

TMS-EEG measures, at scalp electrodes, how perturbation is expressed and propagated across the network by capturing their downstream electrophysiological effects. The effect being confined to specific sites, hemispheres, and frequency bands is what a targeted, system-level recalibration would predict, more than a uniform cortical disinhibition would. Changes in recurrent amplification or in the operating state of the distributed network could produce a similar signature; increased gain is, in our opinion, the most economical of these accounts, and the one that also closes the loop with the SSVEP gradient described above. It would stabilize cortical activity as retinal drive declines, while remaining bounded by how much signal the periphery still transmits.

The enhanced response also occurred within a pre-existing asymmetry of the visual network. Left occipital stimulation produced widespread, mainly contralateral group differences across alpha, beta, and gamma bands, whereas right occipital stimulation produced a more circumscribed, ipsilateral pattern over parietal and central regions, closely matching the asymmetric propagation we have previously described in healthy participants after occipital stimulation (Siviero et al., 2023; Bonfanti et al., 2024; Paolini et al., 2024). RP therefore appears to modulate responsiveness within an already asymmetric architecture. More importantly, the two hemispheres were not affected identically: the early N35 difference was specific to left stimulation, while the more extensive beta-band topological changes followed right stimulation. Averaging across stimulation sides, as is still common practice, would have concealed this, leaving a misleading impression that one hemisphere is simply more responsive than the other.

One further finding deserves consideration: after motor-cortex stimulation, resting motor threshold and the earliest motor TEP components did not differ between groups, and the bilateral occipital beta effect was absent, arguing against a generalized difference in stimulation dose or in cortical responsiveness outside the visual system. Patients did, however, show stronger alpha-band ERSP and altered alpha connectivity after motor-cortex stimulation, which suggests that the consequences of long-standing retinal deafferentation are not confined to occipital cortex. This also means that we cannot attribute every later TMS-EEG difference exclusively to the direct transcranial effect of the pulse: TMS produces auditory and somatosensory responses that can contribute to later components, and the present protocol did not include an optimized sham condition. The early, local N35 effect, the site- and frequency-specific pattern of the later effects, and the absence of any pre-TMS group difference in brain connectivity argue for a genuine cortical contribution, but the motor-cortex finding qualifies how far that contribution can be generalized.

These results carry a specific implication for strategies aimed at restoring retinal input. A weak or absent SSVEP should not be read as evidence that the downstream cortex has become uniformly unresponsive: the central and intermediate visual field still supported measurable entrainment, and direct perturbation uncovered substantial local and network-level capacity to respond. Such preserved, or enhanced, responsiveness could provide a substrate on which restored or newly introduced input can be processed. Greater responsiveness is not, however, necessarily an advantage: if it reflects increased gain, it may amplify noise and aberrant retinal activity as readily as it amplifies a useful signal. Retinal rescue and cortical readaptation may therefore need to be planned together, particularly when a restorative signal differs in strength or statistical structure from the degraded input the cortex has adapted to over the years (Castaldi et al., 2016). We also stress that enhanced TMS-evoked responsiveness is not evidence of preserved conscious visual experience: we did not measure phosphene perception, and how perturbational capacity relates to conscious visual content remains an open question.

Some limitations set the scope of what we can conclude. Our patient sample was small and genetically heterogeneous, which prevents us from separating disease stage from individual variability, and the cross-sectional design cannot establish whether enhanced cortical responsiveness develops progressively with retinal loss or precedes it. SSVEP rings and TMS targets were also not retinotopically matched, so we can describe a system-level dissociation between visually driven activity and perturbational responsiveness, but not whether reduced entrainment in each ring corresponds to altered reactivity in its cortical representation. Longitudinal studies with individualized retinal measures and retinotopic mapping will be needed to establish how the perturbational phenotype we describe relates to disease progression and to the efficacy of visual restoration.

In conclusion, RP does not weaken the visual cortex uniformly: it changes the relationship between residual visual input and the state of the cortical network. Visually driven activity declined from central to peripheral vision and showed sector-specific changes in timing, while direct perturbation of the same cortex elicited enhanced local responses and a stronger, asymmetric distribution of perturbation-related activity through the network. A single mechanism, a homeostatic increase in cortical gain consistent with the plasticity described by Lunghi, Castaldi, Morrone and colleagues in RP and in short-term visual deprivation, offers a simple, but not simplistic, account of both observations at once. Its cellular basis remains to be established, but the evidence strongly supports a broader conclusion: in RP, the deafferented visual cortex retains, and in places strengthens, its capacity to respond as the input that reaches it deteriorates.

